# HBN-EEG: The FAIR implementation of the Healthy Brain Network (HBN) electroencephalography dataset

**DOI:** 10.1101/2024.10.03.615261

**Authors:** Seyed Yahya Shirazi, Alexandre Franco, Maurício Scopel Hoffmann, Nathalia B. Esper, Dung Truong, Arnaud Delorme, Michael P. Milham, Scott Makeig

## Abstract

The Child Mind Institute (CMI) Healthy Brain Network (HBN) project has recorded phenotypic, behavioral, and neuroimaging data from ∼5,000 children and young adults between the ages of 5 and 21. Here, we present HBN-EEG, the “analysis-ready” data from its high-density (128-channel) electroencephalographic (EEG) recording sessions formatted as Brain Imaging Data Structure (BIDS) datasets. HBN-EEG also includes behavioral and task-condition events annotated using Hierarchical Event Descriptors (HED), making the datasets analysis-ready for many purposes without ‘forensic’ search for unreported details. We also ensured data consistency and event integrity and marked inconsistencies. HBN-EEG sessions include six tasks, three with no participant behavioral input (passive tasks) and three including button press responses following task instructions (active tasks). Openly available participant information includes age, gender, and four psychopathology dimensions (internalizing, externalizing, attention, and p-factor) derived from a bifactor model of questionnaire data. Currently, HBN-EEG data from more than 2,600 participants is freely available on NEMAR (nemar.org) and OpenNeuro in the form of nine dataset releases, with further dataset releases to follow.

## Background & Summary

The recent Healthy Brain Network (HBN) project is a large-scale multimodal data collection project of the Child Mind Institute (childmind.org) to collect data from and provide diagnostic insights to children and adolescents (5-21 years) in the greater New York City metropolitan area [1]. HBN project data includes structural and functional MR head imaging, high-density EEG with concurrent eye-tracking and behavioral data, cognitive and psychiatric assessments, and genetic and phenotypic data batteries collected from about 5,000 participants. HBN will continue to collect phenotypical data and other data modalities from this cohort. HBN is collecting novel synchronous experiments [2] that include the following modalities: EEG, eye tracking, physiological measures (respiratory belt, electrocardiogram, electromyography, electrodermal activity), voice and video recordings, actigraphy, body motion tracking, and using tablets for digital tracing. The HBN data are available as an International Neuroimaging Data-sharing Initiative (INDI) resource^1^. The EEG data collection session of the HBN project included ∼60-minute high-density EEG (128 channels; GSN 200, Magnetism-EGI, Eugene, OR) and concurrent eye-tracking recordings. Participants were seated in a dark room, resting their head on a chin rest, viewing a computer screen. During each session, passive tasks (involving no motor responses) included Resting State, Surround Suppression, and Movie Watching (four short films of different types). Active tasks (involving task-based participant mouse click responses) included Sequence Learning, Contrast Change Detection, and Symbol Search. Participants were given a computer mouse with specific instructions for each active task to provide the response using the mouse clicks. Detailed experiment designs and their motivating hypotheses have been detailed by Langer and colleagues [3].

The six tasks were administered by a Linux workstation running Psychtoolbox 3 [4,5], controlling separate workstations recording the EEG and eye-tracking data, respectively. The governing workstation presented audiovisual stimulation and recorded behavioral (mouse click) responses. Photogrammetric scans of participant electrode locations were recorded at the end of the sessions of ∼2,200 participants.

We imported, synchronized, and integrated the behavioral data as event markers in the EEG data files and replaced the numerical event codes used by the experiment control workstation with meaningful descriptive terms. Next, we provided formal annotations describing each of these event markers using Hierarchical Event Descriptors (HED, hedtags.org) [6,7] so that mega-analysis investigations, including machine learning across tasks and datasets can be performed readily and efficiently. We also performed basic “quality checks” on each dataset to select the datasets with available data and event information (Figure 1).

**Figure 1:**
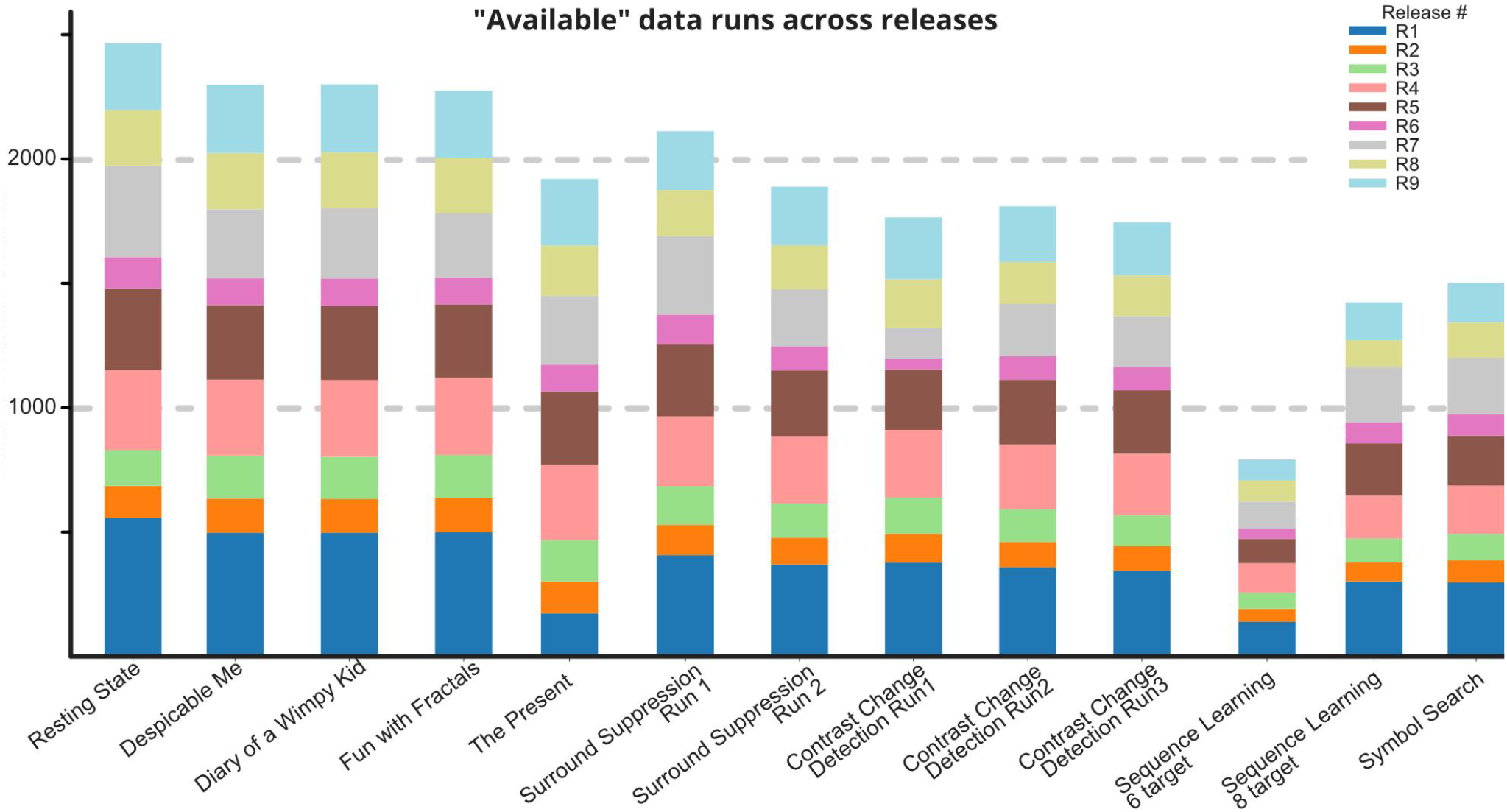
Number of available EEG data runs across the first nine HBN Releases. “Available” flags are determined based on the data length, sampling frequency and availability of event markers (see Methods section).

The initial 9 Releases whose availability is described here include EEG, behavioral, and participant measures from 2639 participants (2634 participants have at least one dataset marked as “available”, Figures 1 and 2 and Table 1). Participants were 35% female, with a median age of 10.38 years, and mainly right-handed, with a median Edinburgh Handedness Questionnaire (EHQ) score of 76.7 [8] (Figure 2). Upon full processing and release, the available HBN-EEG datasets will include data from all ∼5,000 participants, along with eye-tracking and personalized computational head models. The Chesapeake Institutional Review Board approved the study. Researchers obtained written informed consent from participants aged 18 or older, and for those under 18, they obtained consent from legal guardians and permission from the participants. Subjects were anonymized using the Global Unique Identifier (GUID) Tool from the National Institute of Mental Health (NIMH) Data Archive (NDA) [9]. The GUID does not share personally identifiable or protected health information, therefore can be openly shared.

**Table 1.** HBN participant count across all releases. NC indicates subjects with CC-BY-NC-SA4.0 license, which are not included in the OpenNeuro datasets. These subjects are all included in a separate dataset available on an AWS S3 bucket.

| Release Number | Total Participants | Participants > 1 task marked "available" | Participants with bifactor metrics | Digital Object Identifier (DOI) |
| --- | --- | --- | --- | --- |
| R1 | 596 | 578 (136+442NC) | 576 | 10.18112/openneuro.ds005505 |
| R2 | 156 | 152 (148+4NC) | 153 | 10.18112/openneuro.ds005506 |
| R3 | 186 | 183 (182+1NC) | 182 | 10.18112/openneuro.ds005507 |
| R4 | 359 | 324 | 350 | 10.18112/openneuro.ds005508 |
| R5 | 369 | 330 | 359 | 10.18112/openneuro.ds005509 |
| R6 | 143 | 134 | 139 | 10.18112/openneuro.ds005510 |
| R7 | 449 | 381 | 447 | 10.18112/openneuro.ds005511 |
| R8 | 356 | 257 | 356 | 10.18112/openneuro.ds005512 |
| R9 | 364 | 295 | 362 | 10.18112/openneuro.ds005514 |
| R10 | 576 | TBD | 571 | n/a |
| R11 | 571 | TBD | 571 | n/a |
| NC | 0 (already counted) | (447, moved from other datasets) | – | On Amazon S3:<br>s3://fcp-indi/data/Projects/HBN/BID<br>S_EEG/cmi_bids_NC |
| **SUM** | **4125** | **2634** | **4063** |  |

**Figure 2:**
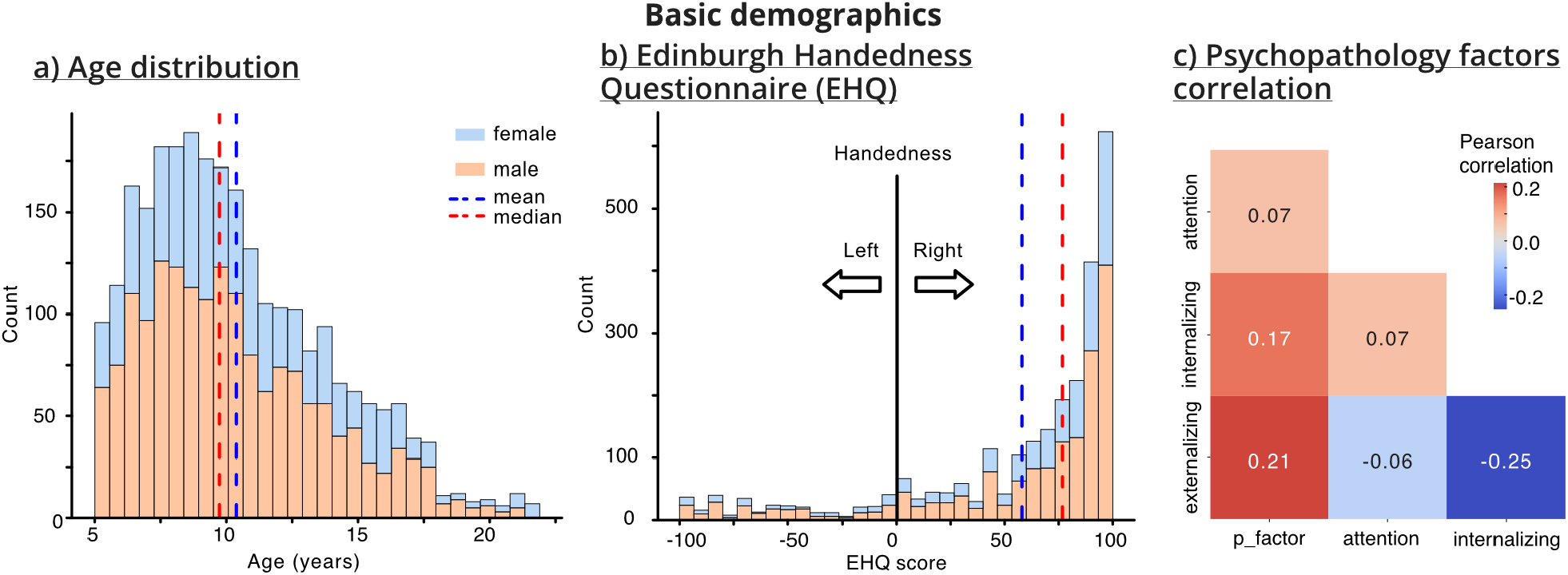
Basic demographics of the HBN-EEG participants across Release 1 to release 9 (2634 subjects), a) Stacked age histogram based on gender, b) Handedness score (−100 indicates left handedness, and 100 indicates right handedness), c) The Pearson correlation between the four psychopathology factors identified for each participants based on their Child Behavior Checklist (CBCL) responses.

Optimal curation of “analysis-ready” datasets is a continuing process. We hope for feedback from the research community to ensure the HBN-EEG dataset is truly “analysis-ready” for a wide variety of research aims. We regard the efforts reported here as a starting point in providing a large, transparent dataset in a form that will assist researchers in easily identifying the information they need to pursue their research without further need to contact the data collection team.

## Methods

While we made our best efforts to preserve the raw nature of the data, several steps were required to make the HBN data “analysis-ready.” The typical archival process for each dataset is detailed in Figure 3. Of note, we first replaced all event codes with more meaningful string values, added subject response and environmental context information from HBN behavioral records, and then annotated all available information concerning sensory, behavioral, and task condition events using the HED system (Generation 3, hedtags.org). We also performed a series of data quality checks to evaluate and ensure data consistency, and integrated available psychopathology factors.

**Figure 3.**
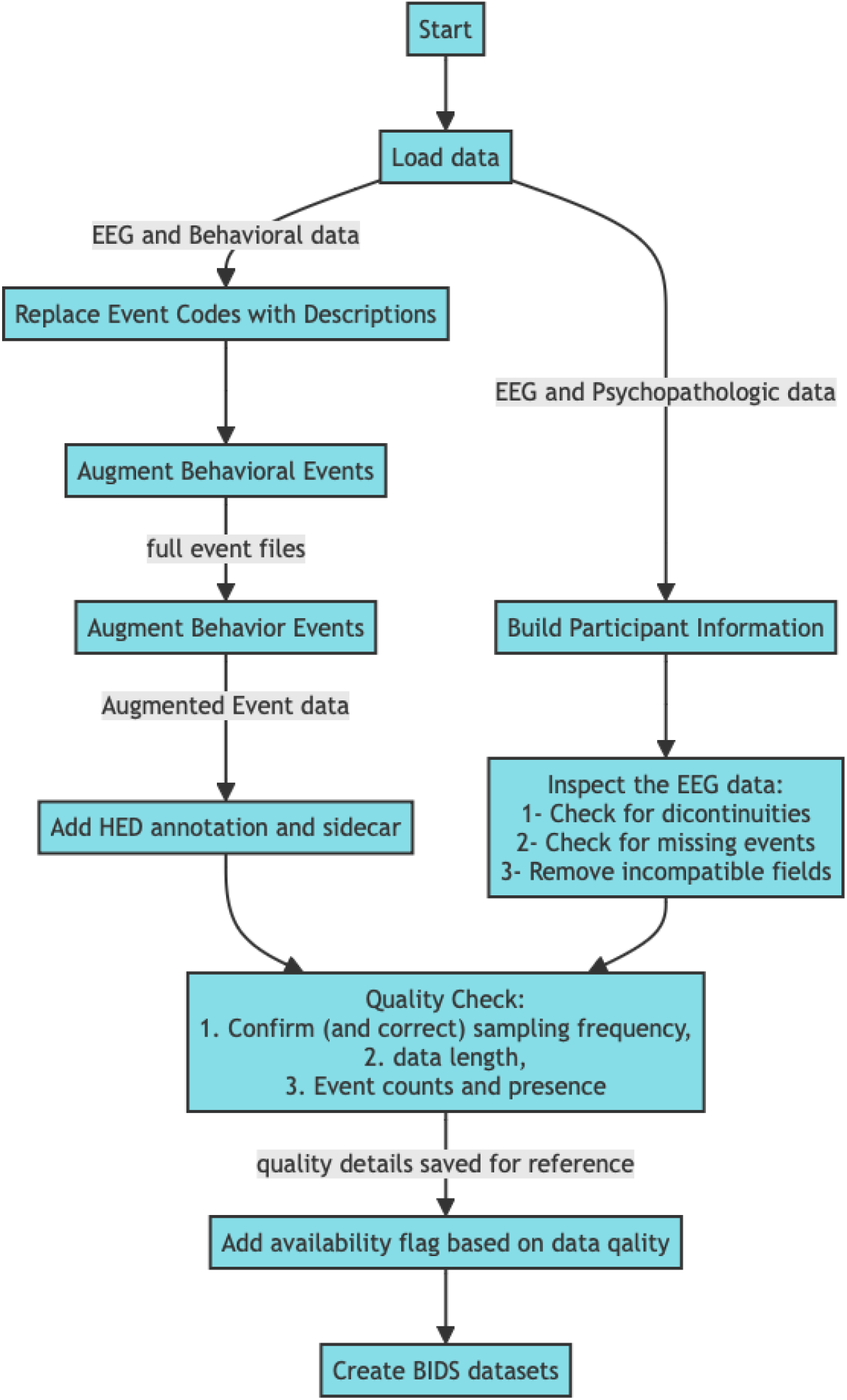
Pipeline for the HBN raw data curation, augmentation and data quality checks

The HBN-EEG datasets, with the publicly available metadata, are now hosted on OpenNeuro and NEMAR in BIDS format to facilitate further data reuse, comparison, or reproduction. The following details of these efforts are given in two sections, namely, whole-session procedural information and task-specific considerations.

### Standard procedures

Our standard process for incorporating additional available data into the dataset and for checking data quality and metadata integrity for each participant involved adding psychopathology factors, testing data and event consistency, adding both required and recommended BIDS metadata, and converting the datasets to the EEG-BIDS format [10].

#### Psychopathology factors

We integrated four available dimensions of psychopathology factors into the HBN-EEG obtained by projection of participant questionnaire responses to the Child Behavior Checklist (CBCL) [11–13]. CBCL is a parent-reported assessment of 120 emotional and behavioral symptoms of the young person (6 to 22 year olds in this study), harmonized with the CBCL Pre (1.5 to 5 year olds, 48.8% CBCL-matched items, 2.4% of the participants), answered in a 3-point scale (0 = not true; 1 = somewhat/sometimes true; 2 = very true/often). For this study, we selected the McElroy et al., 2018 model [11], containing 66 items and four factors, as the model demonstrated to be reproducible in different conditions [12]. The four factors are summarized in the `participants.tsv` file at the root of the (Release) datasets. These factors were derived from a bifactor model that distinguished between general and specific psychological factors (i.e., an orthogonal model), providing a refined perspective on each participant’s mental health. The detailed procedure for obtaining the bifactor model and estimating its validity is described below. These factors can improve the accuracy, statistical rigor, and clinical utility of downstream data analysis compared to the raw questionnaire results [11].

##### P-factor

The p-factor is a general psychopathology dimension computed to estimate psychological common variance across various disorders. An elevated p-factor score indicates heightened general psychopathology, signaling broader mental health risk.

##### Attention

This factor evaluates focus and sustained attention capabilities after the general factor is factored out. Lower attention scores suggest robust attentional control, while higher scores may indicate attentional deficits and hyperactivity, such as those observed in ADHD.

##### Internalizing

The internalizing factor measures symptoms related to distress and fear, such as depression and anxiety, after the general factor is factored out. Higher scores reflect more severe internalizing symptoms, impacting emotional and psychological well-being.

##### Externalizing

Externalizing scores assess issues in outwardly directed behavior, including aggression and rule-breaking, after the general factor is factored out. Higher scores typically denote stronger behavioral challenges and self-regulatory difficulty.

Changes in these scores over time can provide critical clinical insights. For example, a reduction in the p-factor could denote an improvement in overall mental health. In contrast, a rise in specific factors, for example, attention or internalizing, might suggest a developing or worsening condition.

By incorporating these factors into the HBN dataset, we enhance the utility of the dataset for investigating complex interactions between mental health dimensions in young populations, thereby supporting more targeted interventions and research to advance clinical and research methodologies in line with our broader commitment to provide an “analysis-ready” dataset that enables a range of sophisticated and efficient analyses without requiring researchers to design and perform extensive dataset preprocessing.

#### Overview of the Biofactor Model

We built the bifactor model using Confirmatory Factor Analysis (CFA) in Mplus version 8.6 [14]. All 66 CBCL items were configured to load on a general psychopathology factor (the p-factor). Only eight, 31, and 27 items loaded on the attention, internalizing and externalizing, respectively, were loaded for the specific factors. Specific factors were not allowed to correlate with each other, nor with the general factor. CFA was carried out using delta parameterization and weighted least squares with diagonal weight matrix with standard errors and mean- and variance-adjusted chi-square test statistics (WLSMV) estimators. To evaluate model fit, we used root mean square error of approximation (RMSEA), comparative fit index (CFI), Tucker–Lewis index (TLI), and standardized root-mean-square residual (SRMR). Values of RMSEA lower than 0.060 and CFI or TLI values higher than 0.950 indicate a good-to-excellent model. SRMR lower or equal to 0.100 indicates adequate fit and lower than 0.060 in combination with previous indices indicates good fit [15]. These thresholds tend to be less reliable when using categorical data, as in this analysis. The probability of correctly rejecting a misspecified model decreases with higher sample sizes, and CFI/TLI is higher (and RMSEA is lower) when the number of dimensions and response categories increases, and the number of items decreases [16–18]. Therefore, caution is warranted while interpreting fit indexes.

We used nine model-based reliability indices to evaluate the bifactor models. Briefly, they were omega (ω), hierarchical omega (ωH), factor determinacy (FD), H index, explained common variance (ECV) of a specific factor due to itself (ECV-SS), ECV of a specific factor with respect to the general factor (ECV SG), ECV of the general factor with respect to a specific factor (ECV GS), percent uncontaminated correlations (PUC) and item explained common variance (IECV) [19,20]. When ωH is > 0.8 and ECV and PUC are > 0.7, the construct can be interpreted as unidimensional [20]. All bifactor reliability indices were calculated using the BifactorIndicesCalculator package in R [19].

The bifactor model demonstrated an acceptable fit to the data (RMSEA = 0.052, 95% CI [0.052, 0.053], CFI = 0.844, TLI = 0.833 and SRMR = 0.076). Model-based indices demonstrated good reliability for the factors, with high FD and H indexes for all factors. IECV demonstrated that symptoms of crying, preferring to be with older people, setting fire, thinking about sex, stubbornness, mood changes, sulking and getting suspicious are stronger indicators of the general factor. General model structure with factor loadings and model-based reliability indices can be found in Supplementary Table S1.

Researchers can access additional phenotypic data from the HBN data web portal after providing their rationale and signing a data use agreement. Additional available domains include data on participant (a) demographics, (b) cognition,(c)language, (d) emotional and psychological function, (d) social, (e) emotional, and (f) behavioral functioning, (g) family structure, (h) stress and trauma, (i) physical skills, (j) fitness, and (k) physiologic status of substance use and addiction, (l) neurologic function, (m) eye movements, and (n) medical status and diagnosis.

#### Data and event consistency assurance

Ensuring the individual data runs (i.e., data recordings) load and have appropriate features, including data sampling rate, event stream, and duration, is critical for any successful downstream use of the data. Without such quality assurance, each researcher would need to develop custom data ingestion and curation pipelines to exclude unfit data. Here, we introduce “availability” flags for each data run in the `participants.tsv` file so researchers can easily filter through the data runs and only choose the available data runs. Data availability flags include **Available, Caution**, and **Unavailable**. Available data runs are guaranteed to load and include timings and descriptions of key events; data runs with Caution flags load successfully but fail at least one consistency test. Unavailable data runs are not included in the dataset. The tests we ran on each data run included (1) when needed resampling data runs to 500 Hz, (2) checking for abnormal data run duration, (3) confirming the presence of key task events (e.g., task start and stop), and (4) checking for abnormal length of the recorded events. We labeled data runs outside five times the interquartile range (IQR) of recorded event length or data run duration with a ‘Caution’ flag. The detailed results of these quantity assurance tests per task are available as tables in the *code/* directory of each Release dataset.

#### BIDS metadata

BIDS provides a structure for data file names and formats, and a suite of required and recommended metadata fields essential for reproducing or performing analysis of the datasets [10,21]. We used the original experiment design manuscripts [1,3], cross-checked with the experiment implementation scripts provided by the Child Mind Institute and the HBN data portal^2^ to complete the dataset-level and task-level metadata.

### Task-specific procedures

Modifications specific to each task include importing behavioral responses and environment conditions for the individual EEG data runs, and constructing HED annotations for the event files. The information added for the passive (Resting State, Surround Suppression, and Movie Watching) and active (Contrast Change Detection, Sequence Learning, and Symbol Search) tasks are listed below. Detailed descriptions of each task are available in [3]. Below, we highlight the specific annotation enhancements we performed.

#### Resting State

In the Resting State task, participants rested their heads on a chin rest, looking forward to the computer screen with a fixation cross at the center of the screen. A pre-recorded voice instructed them to hold their eyes closed or open and fixate on the central cross during ‘eyes open’. The timeline of the event marker for the task is shown in Figure 4.

**Figure 4:**
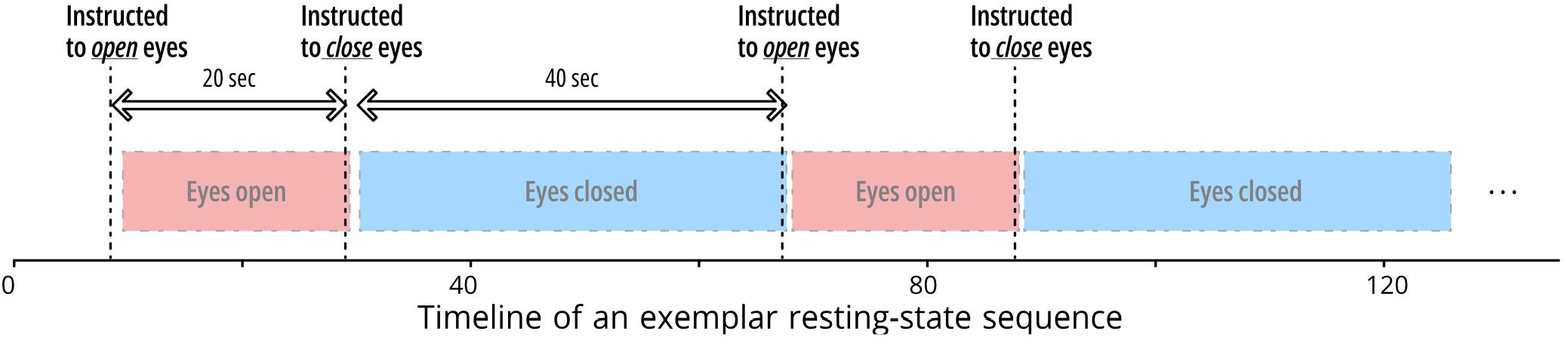
The resting-state instruction and event timeline. The exact eyes open and closed time should be determined using eye-tracking data (will be shared in BIDS format soon)

Each HBN-EEG task includes detailed HED descriptions of event markers; these provide machine-readable and human-understandable event details suitable for a wide variety of analyses. For example, the HED description of the pre-recorded voice instructing subjects to open their eyes during the Resting State task is:

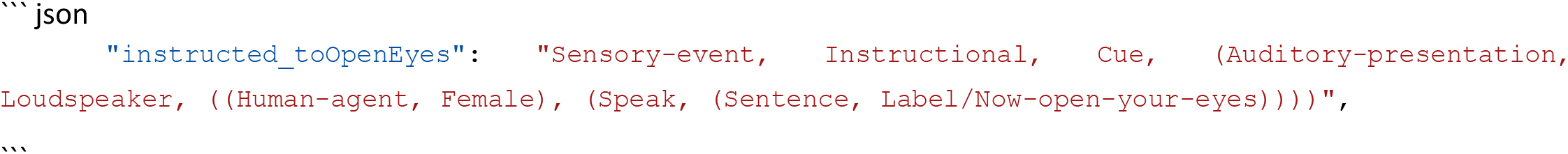

From the description, we can identify that the pre-recorded instruction was delivered using a female voice as a complete sentence (“Now open your eyes.”), giving an instructional cue. Similar details for all HBN-EEG tasks are available in the datasets and Supplements. The HED annotation for the Resting State task is included in the Supplementary material S2.

#### Surround Suppression

Participants observed a modified Surround Suppression stimulus sequence (derived from [22]). Four flashing peripheral disks in the foreground were presented with a contrasting background in two runs, each ∼3.6 minutes long. We imported and synchronized foreground and background stimulus parameters from the corresponding behavioral recording for each subject to complete the stimulus event description. Figure 5 shows an example of an actual event-marker stream during the Surround Suppression task. The HED annotation of the event markers is available in Supplementary material S3. The stimulus representations and the event timeline in Figure 5 were drawn using the codes generated by a Large Language Model (Claud Sonnet 3.5 [23]) directly from the HED stimulus descriptions with minor final edits.

**Figure 5:**
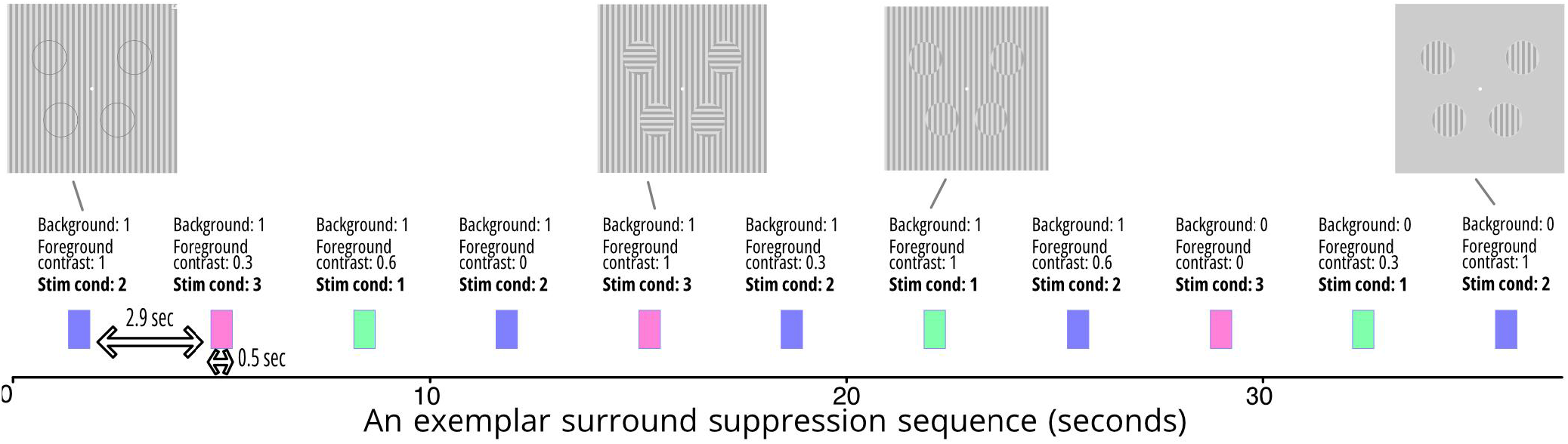
An exemplar surround suppression even stream. There are three factors modulating stimulus presentation, stimuli condition (noted as Stim cond in the image, that is, the relation between foreground and background), foreground contrast, and background presence. Here, the colored event markers indicate different stimuli conditions.The foreground disks flicker on and off at 25 Hz. The exemplar stimuli visual presentations depict an estimate of the static image of the stimuli.

#### Movie Watching

The final passive tasks involved watching movies. Participants were asked to watch four short movies with different themes and aims (1) ‘Despicable Me’, (2) ‘Diary of a Wimpy Kid’, (3) ‘Fun with Fractals’, and (4) ‘The Present’ [1]. Details and rationale for the movie choice and links to access the presented movies are available in the top-level _eeg.json files. The recorded event markers for each movie presentation include only the start and stop times. We will soon introduce methods and remodeling tools to insert custom event sequences for each movie relative to these event anchors (movie presentation start and stop) so researchers will be able to plug in an event stream of choice. We will also provide an example detailed annotation in HED format for the animated drama ‘The Present.’

#### Contrast Change Detection

In this task, two co-centric flickering grated disks (one with left-leaning gratings and one with right-leaning gratings) appear on the screen. After a randomly selected time, one of the two flickering grated disks on the screen would change the contrast. Participants were asked to identify the left-leaning or right-leaning grated disks with dominant contrast as soon as they could identify the disk. Based on their responses, feedback (a smiley face or sad face) was presented. We imported participant responses from the behavioral recordings and assembled HED descriptions for the main task and the respective subject responses. An example event marker sequence is shown schematically in Figure 6. HED event descriptions are available in Supplementary material S4.

**Figure 6.**
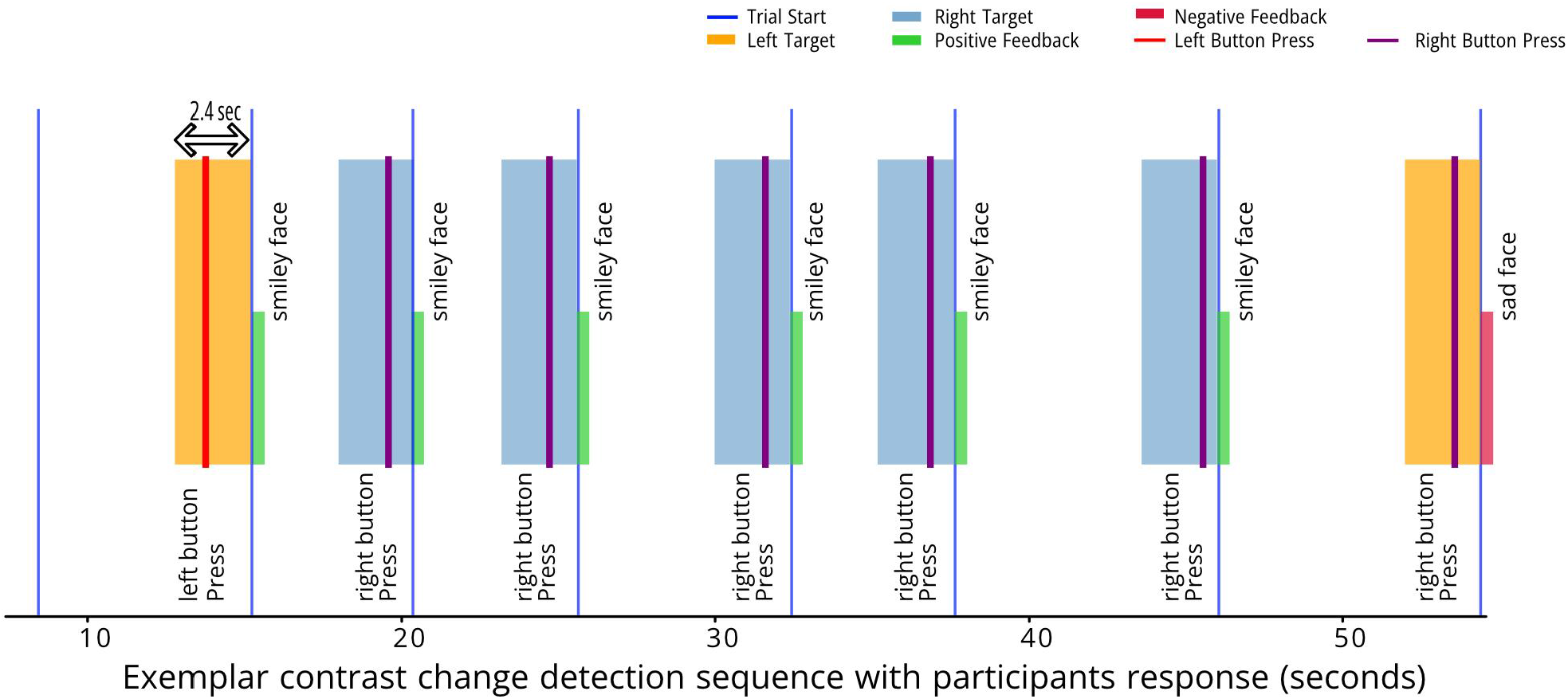
Exemplar timeline of a few contrast change detection stimulus presentations. The left- and right leaning disks were flashing at different frequencies while one disk would increase contrast and rapidly resetting to the original contrast levels. The participant had to provide left or right button press indicating the perceived dominating contrast. The feedback were presented as an smiley or sad face after then of each stimulus presentation.

#### Sequence learning

Participants were shown a sequence of 10 (or 7 if <8 years old) flashed circles among 8 (or 6) possible and predetermined targets positioned on the periphery of a larger invisible circle on the screen (Figure 7). The same sequence was repeated five times, and after each repetition, participants were asked to repeat the sequence using a computer mouse. The participant’s response sequence was recorded for each repetition, but unfortunately, the timestamps of this response were not recorded. Because of the difference in the target and sequence counts, we divided this task into two tasks: a six-target task, and an eight-target task. The task is based on a similarly designed sequence learning experiment by Steinemann [24]. An exemplar timeline of the sequence presentation block is presented in Figure 7 and annotated using HED tags in Supplementary material S5 (for the eight-target task).

**Figure 7.**
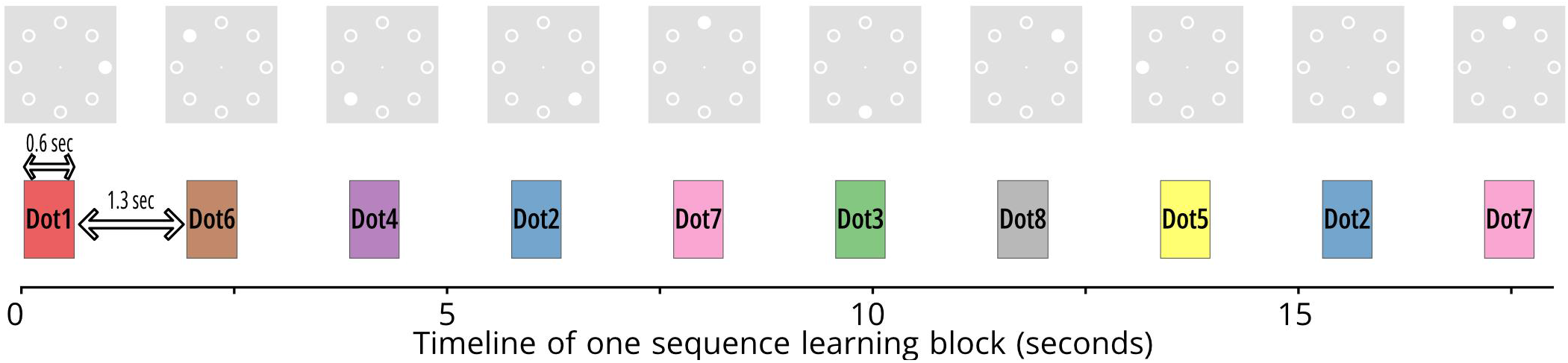
The stimuli timeline for the sequence learning task with 10 targets for participants older than 8 years old.The exemplar stimuli visual presentations depict an estimate of the static image of the stimuli. See the stimuli screen casts for the actual stimuli presentation.

#### Symbol search

The final task in the interactive battery of experiments involves an emulation of a standard neuropsychological test (a subset of the Wechsler Intelligence Scale for Children IV, WISC-IV [25]) on the computer screen, in which participants were asked to confirm if either of the target symbols in each row were present in the five search symbols present in the same row (Figure 8). The test was presented to the participants in 15-row sequences, and participants were asked to go to the next set of sequences once they responded to all 15 rows. Since each of the 15 questions was presented simultaneously, only the times participants pressed the mouse button to respond were recorded. The HED description for the event makers of this task is in Supplementary material S6.

**Figure 8:**
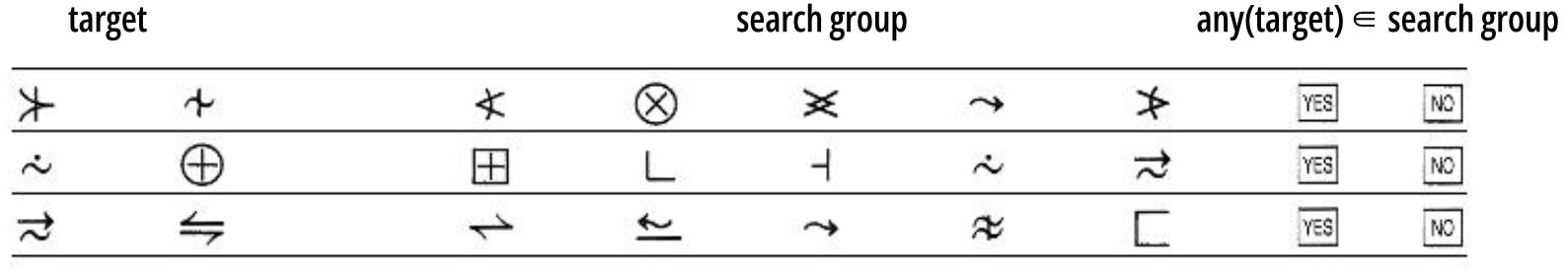
Exemplar 3 questions of the sequence learning task. Participants were instructed to click on Yes if any of the target shapes were present in the search group.

## Data Records

The HBN project currently has 11 Releases with over 4000 participants (Table 1). There will be an additional release that will conclude the data release of this project with the current EEG protocol (in total, 12 Releases). Our first dissemination of HBN-EEG will include the first nine Releases as one BIDS dataset per Release (nine total BIDS datasets) [26–34]. We did not include Release 10 and 11 because many of the individual data files currently lack the metadata needed to assemble the BIDS datasets. Fortunately, the psychopathology factors have been quantified using all 12 Releases, so we do not expect the participant information to change in the subsequent releases (except for possible increases in data availability).

The datasets are available under two licenses, CC-BY-SA 4.0 and CC-BY-**NC**-SA 4.0, based on the license consented by the participants. We only included the datasets under CC-BY-SA 4.0 in the nine Releases hosted on OpenNeuro. A single dataset containing all participants with CC-BY-NC-SA 4.0 is available from the FCP-INDI Amazon Web Services (AWS) S3 bucket (Table 1). All datasets are also available on NEMAR with proper license attribution.

### Future development

Apart from adding more participants, we are planning to complement HBN-EEG datasets with the following data:

1. **Eye-tracking:** EEG tasks were performed in conjunction with eye-tracking using table-top eye trackers. We have already developed the remodeling tools to convert the eye-tracking data to BIDS. We will update all datasets with eye-tracking information as soon as the Eyetracking-BIDS specifications are finalized.
2. **Electrode locations:** Around 2270 of the HBN participants also have EEG electrode locations scanned and ready for electrode localization (i.e., electrode digitization). We are developing an open-source automated electrode localization toolbox to objectively digitize the electrodes.
3. **High-density leadfield matrix:** There has been a significant effort to use the structural MRI scans of each HBN participant to create personalized computational head models and lead-field matrix for EEG source imaging [35]. The personal lead-field matrix will help determine the sources of EEG signals with centimeter accuracy. The Riaz and colleagues’ effort creates lead-field matrices with ∼8000 nodes within the cortical volume with uniform distribution. We will extend the lead-field matrix resolution by ten fold to ∼80,000 nodes, which also takes the cortical surface normals into effect using the NFT toolbox [36]. This step is essential for high-resolution EEG source imaging, especially for developing children and adolescents [37,38].
4. **Skull conductivity estimation:** Skull to Brain conductivity ratio plays a crucial role in the accuracy of EEG source models [39]. Skull conductivity decreases about 10-fold from adolescence to 50 years of age and is largely varied across subjects and measurement methods [40–42]. Therefore, we will use the simultaneous tissue Conductivity And source Location Estimation (SCALE) method to estimate the brain-to-skull conductivity ratio (BSCR) for each participant and will regenerate and share a new lead field matrix for each individual with the updated values.

## Using HED for transparent experiment annotation

There are three main objectives for using HED for annotating datasets: 1) building event contexts, 2) creating machine-readable and human-understandable annotations for mega-analysis and machine learning, and 3) task transparency and reproducibility. A goal of HED is to provide context for the events happening during an experiment by carefully creating relations between concurrent and precedent events and with as much detail as the researcher has provided [7].

HED also creates a platform for mega- and meta-analysis, in which common event markers can be pulled across studies and brain-imaging modalities to create inferences across studies and find common dynamics for common events and event contexts. HED tags used in the annotations are hierarchical, e.g., the “Press” tag for mouse-button press conveys many more meanings based on the higher nodes in the hierarchy: `Action/Move/Move-body-part/Move-upper-extremity/Press`. So, analysis can be done for the leaf node (in this example, Press) and across all other nodes. This is particularly beneficial when using large machine learning architectures with large (multimodal) datasets to extract manifolds and subspaces that each node shares across datasets and modalities.

The last use case of HED is to provide a common language to describe what happened in the experiment, such as the visual stimulus that the participant observed or the temporal sequence of the experiment events. Since HED is a tightly structured descriptor, Language Models can adequately “translate” HED annotations into other languages. For example, we used Anthropic Claude Sonnet 3.5 to translate the visual presentation of the Surround Suppression task from HED descriptions to Scalable Vector Graphics (SVG) format (Figure 9). The resulting figure accurately represented the stimulus up to simple rotation of foreground circle placements, which was not provided in the HED description in the first place. Similarly, Figures 4 through 7 were all created using Sonnet 3.5 by providing a snippet of the events file and the HED description for the events files. We see HED as a common space that can be translated to different experiment control software or inference algorithms while ensuring that experiment details and contexts are clear and accurate up to the details provided in the HED annotations.

**Figure 9:**
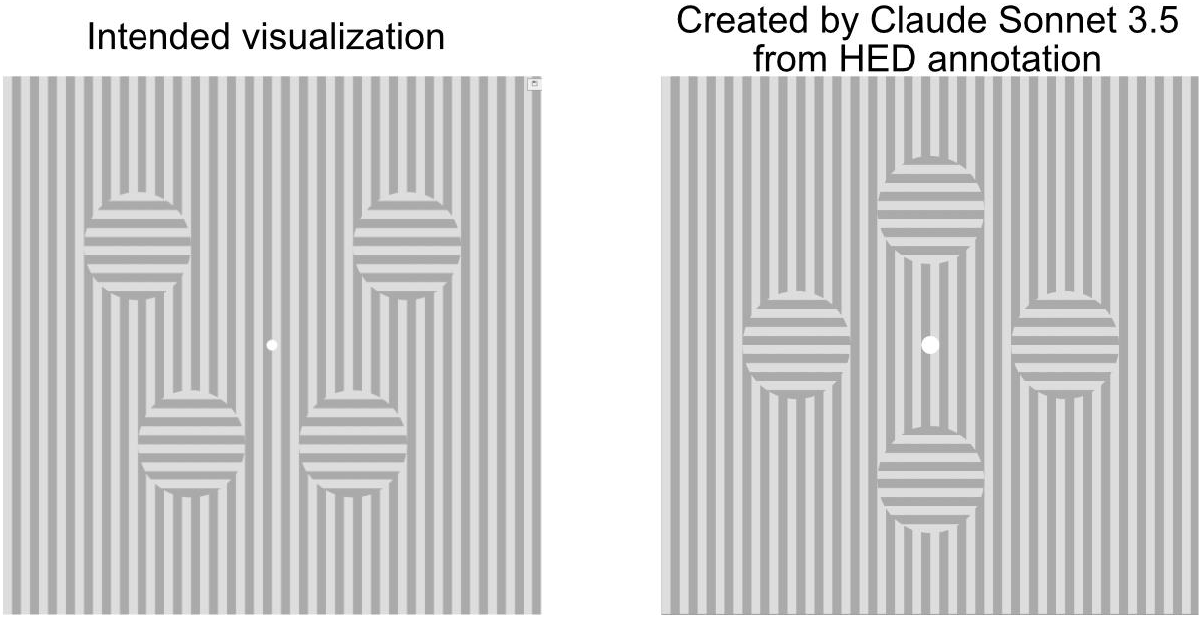
The surround suppression visualization shown to the participant. Left panel is the intended visualization, and the right panel is re-generation of the visualization by Claude Sonnet 3.5 using only the HED annotation.

## Data Availability

All datasets are available on NEMAR and FCP-INDI AWS storage at `s3://fcp-indi/data/Projects/HBN/BIDS_EEG/`. The datasets with a CC-BY-SA 4.0 license are also available on OpenNeuro.

## Acknowledgments

The authors thank Kay Robbins for their invaluable abstinence in composing the HED descriptions.

This work was partially supported by NIH grants R01MH125934, R24MH120037, R01MH120482, and R01NS047293. All data curation and processing have been performed on the Neuroscience Gateway [43,44].

## Author Contributions

SYS curated the datasets, performed quality checks, made the HED annotation, and created the manuscript figures. SYS, AF, AD, and SM wrote and edited the manuscript. MSH quantified the bifactor model and extracted the psychopathology dimensions from this model. AF, AD, DT, and SM provided feedback for data curation. DT helped with HED annotation. AF and NBE managed and retrieved the raw data. AD, SM, and MPM provided funding. MPM conceived the HBN project.

## Footnotes

1 http://fcon_1000.projects.nitrc.org/indi/cmi_healthy_brain_network/sharing_neuro.html

2 https://fcon_1000.projects.nitrc.org/indi/cmi_healthy_brain_network/MRI_EEG.html

