## Supplementary Table S1 for "HBN-EEG: The FAIR implementation of the Healthy Brain Network (HBN) electroencephalography dataset"

Table S1 - HBN CBCL bifactor model (according to McElroy et al. 2018)

| Content | Item | Factors and factor loadings | | | | IECV |
|---|---|---|---|---|---|---|
| | | P-factor | Internalizing | Externalizing | Attention | |
| Cries | CBCL_14 | 0.559 | 0.153 | | | 0.930 |
| Fears | CBCL_29 | 0.336 | 0.342 | | | 0.491 |
| Fears school | CBCL_30 | 0.368 | 0.399 | | | 0.460 |
| Fears do bad | CBCL_31 | 0.446 | 0.341 | | | 0.631 |
| Perfect | CBCL_32 | 0.221 | 0.440 | | | 0.201 |
| Unloved | CBCL_33 | 0.626 | 0.213 | | | 0.896 |
| Worthless | CBCL_35 | 0.508 | 0.460 | | | 0.549 |
| Nervous | CBCL_45 | 0.554 | 0.498 | | | 0.553 |
| Fearful | CBCL_50 | 0.396 | 0.684 | | | 0.251 |
| Feels too guilty | CBCL_52 | 0.317 | 0.580 | | | 0.230 |
| Self-conscious | CBCL_71 | 0.412 | 0.490 | | | 0.414 |
| Talks about suicide | CBCL_91 | 0.569 | 0.194 | | | 0.896 |
| Worries | CBCL_112 | 0.356 | 0.663 | | | 0.224 |
| Prefers alone | CBCL_42 | 0.281 | 0.463 | | | 0.269 |
| Won’t talk | CBCL_65 | 0.450 | 0.291 | | | 0.705 |
| Secretive | CBCL_69 | 0.491 | 0.308 | | | 0.718 |
| Shy | CBCL_75 | 0.184 | 0.504 | | | 0.118 |
| Lacks energy | CBCL_102 | 0.335 | 0.497 | | | 0.312 |
| Sad | CBCL_103 | 0.558 | 0.491 | | | 0.564 |
| Withdrawn | CBCL_111 | 0.335 | 0.557 | | | 0.266 |
| Nightmares | CBCL_47 | 0.372 | 0.244 | | | 0.699 |
| Constipate | CBCL_49 | 0.266 | 0.245 | | | 0.541 |
| Dizzy | CBCL_51 | 0.301 | 0.592 | | | 0.205 |
| Tired | CBCL_54 | 0.406 | 0.468 | | | 0.429 |
| Aches | CBCL_56A | 0.310 | 0.459 | | | 0.313 |
| Headaches | CBCL_56B | 0.252 | 0.555 | | | 0.171 |
| Nausea | CBCL_56C | 0.274 | 0.730 | | | 0.123 |
| Eye problems | CBCL_56D | 0.165 | 0.241 | | | 0.319 |
| Skin problems | CBCL_56E | 0.237 | 0.232 | | | 0.511 |
| Stomach | CBCL_56F | 0.271 | 0.621 | | | 0.160 |
| Vomit | CBCL_56G | 0.264 | 0.407 | | | 0.296 |
| No guilt | CBCL_26 | 0.593 | | 0.418 | | 0.668 |
| Bad friends | CBCL_39 | 0.496 | | 0.311 | | 0.718 |
| Lies or cheats | CBCL_43 | 0.598 | | 0.359 | | 0.735 |
| Prefers older | CBCL_63 | 0.409 | | 0.099 | | 0.945 |
| Sets fires | CBCL_72 | 0.444 | | 0.158 | | 0.888 |
| Steals from home | CBCL_81 | 0.492 | | 0.527 | | 0.466 |
| Steals outside home | CBCL_82 | 0.424 | | 0.559 | | 0.365 |
| Swears | CBCL_90 | 0.561 | | 0.261 | | 0.822 |
| Thinks about sex | CBCL_96 | 0.547 | | 0.098 | | 0.969 |
| Argues | CBCL_3 | 0.709 | | 0.331 | | 0.821 |
| Mean | CBCL_16 | 0.591 | | 0.528 | | 0.556 |

| | | | | | | |
|---|---|---|---|---|---|---|
| Demands a lot of attention | CBCL_19 | 0.638 | | 0.233 | | 0.882 |
| Destroys own things | CBCL_20 | 0.530 | | 0.590 | | 0.447 |
| Destroys other | CBCL_21 | 0.528 | | 0.691 | | 0.369 |
| Disobedient at home | CBCL_22 | 0.699 | | 0.494 | | 0.667 |
| Disobedient at school | CBCL_23 | 0.515 | | 0.463 | | 0.553 |
| Fights | CBCL_37 | 0.591 | | 0.487 | | 0.596 |
| Attacks | CBCL_57 | 0.534 | | 0.577 | | 0.461 |
| Screams | CBCL_68 | 0.671 | | 0.323 | | 0.812 |
| Stubborn | CBCL_86 | 0.796 | | 0.145 | | 0.968 |
| Mood changes | CBCL_87 | 0.809 | | -0.044 | | 0.997 |
| Sulks | CBCL_88 | 0.832 | | -0.209 | | 0.941 |
| Suspicious | CBCL_89 | 0.746 | | 0.013 | | 1.000 |
| Teases | CBCL_94 | 0.588 | | 0.366 | | 0.721 |
| Temper | CBCL_95 | 0.736 | | 0.340 | | 0.824 |
| Threatens | CBCL_97 | 0.626 | | 0.539 | | 0.574 |
| Loud | CBCL_104 | 0.584 | | 0.234 | | 0.862 |
| Acts too young | CBCL_1 | 0.419 | | | 0.277 | 0.696 |
| Can't concentrate | CBCL_8 | 0.462 | | | 0.697 | 0.305 |
| Can't sit still | CBCL_10 | 0.483 | | | 0.420 | 0.569 |
| Confused | CBCL_13 | 0.478 | | | 0.528 | 0.450 |
| Daydreams | CBCL_17 | 0.349 | | | 0.596 | 0.255 |
| Impulsive | CBCL_41 | 0.742 | | | 0.224 | 0.916 |
| Poor school work | CBCL_61 | 0.431 | | | 0.336 | 0.622 |
| Stares | CBCL_80 | 0.418 | | | 0.546 | 0.370 |
| Index | | | | | | |
| PUC | | 0.607 | | | | |
| ECV SS | | 0.572 | 0.582 | 0.291 | 0.493 | |
| ECV SG | | 0.572 | 0.222 | 0.143 | 0.063 | |
| ECV GS | | 0.572 | 0.418 | 0.709 | 0.507 | |
| Omega | | 0.948 | 0.894 | 0.930 | 0.782 | |
| OmegaH | | 0.750 | 0.502 | 0.219 | 0.374 | |
| H | | 0.966 | 0.905 | 0.853 | 0.736 | |
| FD | | 0.971 | 0.948 | 0.925 | 0.882 | |

Note: IECV, item explained common variance ($\geq$ 0.85 yield unidimensional item sets that reflect the content of the general factor); ECV, explained common variance; SS, proportion of common variance of the items in each factor which is due to that factor; SG, ECV proportion of common variance of the items in each specific factor which is due to the specific factor; GS, proportion of common variance of the items in each specific factor which is due to the general factor; PUC, percent of uncontaminated correlations; OmegaH, omega-hierarchical; H, index of construct replicability (>0.8 suggests a well-defined latent variable); FD, factor determinancy (>0.9 indicate that the factor score can be used)
