## Supplementary material S2 for "HBN-EEG: The FAIR implementation of the Healthy Brain Network (HBN) electroencephalography dataset"

```
{
  "onset": {
    "LongName": "Event onset",
    "Description": "Position (latency) of event marker in seconds relative to the start.",
    "Units": "seconds"
  },
  "duration": {
    "Description": "Duration of the event, set typically to n/a. Instead, the duration of the event is determined by the onset/offset flags or the next event.",
    "Units": "seconds"
  },
  "sample": {
    "Description": "Temporal position (latency) of event marker relative to the start, in data frames."
  },
  "value": {
    "Description": "The event marker value (type).",
    "Levels": {
      "instructed_toOpenEyes": "A voice prompt instructed subject to open their eyes",
      "instructed_toCloseEyes": "A voice prompt instructed subject to open their eyes",
      "resting_start": "Recording of Resting State task started"
    },
    "HED": {
      "instructed_toOpenEyes": "Sensory-event, Instructional, Cue, (Auditory-presentation, Loudspeaker, ((Human-agent, Female), (Speak, (Sentence, Label/now-open-your-eyes))))",
      "instructed_toCloseEyes": "Sensory-event, Instructional, Cue, (Auditory-presentation, Loudspeaker, ((Human-agent, Female), (Speak, (Sentence, Label/now-close-your-eyes))))",
      "resting_start": "(Def/resting-start, Onset)"
    }
  },
  "event_code": {
    "Description": "The original code used during data collection to indicate an event marker.",
    "Levels": {
      "20": "A voice prompt instructed subject to open their eyes",
      "30": "A voice prompt instructed subject to open their eyes",
      "90": "Recording of Resting State task started"
    }
  },
  "hed_definitions": {
    "Description": "",
```

```
    "HED": {
      "resting_state_def": "(Definition/resting-start, ((Visual-presentation, (Foreground-view, (Black, Cross), (Center-of, Computer-screen)), (Background-view, White)), Recording))"
  }
  }
}
```
