## Supplementary material S3 for "HBN-EEG: The FAIR implementation of the Healthy Brain Network (HBN) electroencephalography dataset"

```json
{
  "onset": {
    "LongName": "Event onset",
    "Description": "Position (latency) of event marker in seconds relative to the start.",
    "Units": "seconds"
  },
  "duration": {
    "Description": "Duration of the event, set typically to n/a. Instead, the duration of the event is determined by the onset/offset flags or the next event.",
    "Units": "seconds",
    "HED": "Duration/#"
  },
  "sample": {
    "Description": "Temporal position (latency) of event marker relative to the start, in data frames."
  },
  "value": {
    "Description": "The event marker value (type).",
    "Levels": {
      "surroundSuppB1_start": "Start of surround suppresstion block (run) 1",
      "surroundSuppB2_start": "Start of surround suppresstion block (run) 2",
      "fixpoint_ON": "Fixation point appears on the screen",
      "stim_ON": "Stimulation appears on the screen"
  },
  "HED": {
    "surroundSuppB1_start": "(Def/surroundSup-start, Onset)",
    "surroundSuppB2_start": "(Def/surroundSup-start, Onset)",
    "fixpoint_ON": "(Def/fixpoint, Onset)",
    "stim_ON": "({duration}, (Visual-presentation, (Background-view, {background}), (Foreground-view, (Peripheral-view, (Circle, Item-count/4, (Radius, Angle/2 degree)))), (Luminance-contrast, {foreground_contrast}), (Spatial-relation, {stimulus_cond})))"
  }
},
"event_code": {
"Description": "The original code used during data collection to indicate an event marker",
  "Levels": {
     "93": "Start of surround suppresstion block (run) 1",
     "97": "Start of surround suppresstion block (run) 2",
     "4": "Fixation point appears on the screen",
     "8": "Stimulation appears on the screen"
  }
},
```

```
"background": {
    "Description": "Whether the background is present",
    "Levels": {
      "0": "Background is not present",
      "1": "Background is present, sinusoidal luminance-modulated gratings with a spatial frequency of 1 cycle per degree in all conditions"
    },
    "HED": {
      "0": "(Visual-presentation, (Background-view, Gray))",
      "1": "(Visual-presentation, (Background-view, (Pattern, Vertically-oriented, Label/sinusoidal-luminance-modulated-gratings-with-a-spatial-frequency-of-1-cycle-per-degree-in-all-conditions)))"
    }
  },
  "foreground_contrast": {
    "Description": "The foreground contrast",
    "Levels": {
      "0": "No contrast",
      "0.33": "1/3 contrast",
      "0.66": "2/3 contrast",
      "1": "Full contrast"
    },
    "HED": {
      "0": "(Luminance-contrast, Fraction/0)",
      "0.3": "(Luminance-contrast, Fraction/0.33)",
      "0.6": "(Luminance-contrast, Fraction/0.66)",
      "1": "(Luminance-contrast, Fraction/1)"
    }
  },
  "stimulus_cond": {
    "Description": "The condition of the stimulus",
    "Levels": {
      "1": "Stimulus condition 1: peripheral foreground disks with parallel, spatially opposite-phase background",
      "2": "Stimulus condition 2: peripheral foreground disks with parallel, spatially in-phase background",
      "3": "Stimulus condition 3: peripheral foreground disks with orthogonal background"
    },
    "HED": {
      "1": "(Spatial-relation, Vertically-oriented, Label/peripheral-foreground-disks-with-parallel-spatially-opposite-phase-background)",
      "2": "(Spatial-relation, Vertically-oriented, Label/peripheral-foreground-disks-with-parallel-spatially-in-phase-background)",
      "3": "(Spatial-relation, Horizontally-oriented, Label/peripheral-foreground-disks-with-orthogonal-background)"
```

```
    }
  },
  "hed_defs":{
    "Description": "Metadata dictionary for defining the stimulus",
    "HED":{
      "surroundSup-start_def": "(Definition/surroundSup-start,((Visual-presentation, (Background-view, Gray)), Recording))",
      "fixpoint_def": "(Definition/fixpoint, (Visual-presentation, (Foreground-view, (Circle, Item-count/1, White, (Center-of, Computer-screen)))))",
      "stim_def": "(Definition/stim, (Visual-presentation, (Background-view, Gray), (Foreground-view, (Dots, Item-count/1, White, (Center-of, Computer-screen)))))"
    }
  }
}
```
