## Supplementary material S4 for "HBN-EEG: The FAIR implementation of the Healthy Brain Network (HBN) electroencephalography dataset"

```json
{
  "onset": {
    "LongName": "Event onset",
    "Description": "Position (latency) of event marker in seconds relative to the start.",
    "Units": "seconds"
  },
  "duration": {
    "Description": "Duration of the event, set typically to n/a. Instead, the duration of the event is determined by the onset/offset flags or the next event.",
    "Units": "seconds"
  },
  "sample": {
    "Description": "Temporal position (latency) of event marker relative to the start, in data frames."
  },
  "value": {
    "Description": "The event marker value (type).",
    "Levels": {
        "contrastChangeB1_start": "Start of contrast change detection block (run) 1",
        "contrastChangeB2_start": "Start of contrast change detection block (run) 2",
        "contrastChangeB3_start": "Start of contrast change detection block (run) 3",
        "contrastTrial_start": "Start of a single contrast change trial",
        "right_target": "The right-side leaning target was presented",
        "left_target": "The left-side leaning target was presented",
        "right_buttonPress": "The right mouse button was pressed",
        "left_buttonPress": "The left mouse button was pressed"
    },
    "HED": {
      "contrastChangeB1_start": "(Def/contrastChange, Onset)",
      "contrastChangeB2_start": "(Def/contrastChange, Onset)",
      "contrastChangeB3_start": "(Def/contrastChange, Onset)",
      "right_target": "((Def/right-target), Duration/1600 ms), (Delay/1600 ms, (Def/right-comeback), Duration/800 ms), (Delay/2400 ms, ({feedback}), Duration/400 ms)",
      "left_target": "((Def/left-target), Duration/1600 ms), (Delay/1600 ms, (Def/left-comeback), Duration/800 ms), (Delay/2400 ms, ({feedback}), Duration/400 ms)",
      "right_buttonPress": "(Agent-Action, (Press, (Right, Mouse-button), (Right, Index-finger)))",
      "left_buttonPress": "(Agent-Action, (Press, (Left, Mouse-button), (Left, Index-finger)))"
    }
  },
  "event_code": {
    "Description": "The original code used during data collection to indicate an
```

```
event marker.",
    "Levels": {
        "94": "Start of contrast change detection block (run) 1",
        "95": "Start of contrast change detection block (run) 2",
        "96": "Start of contrast change detection block (run) 3",
        "5": "Start of a single contrast change trial",
        "9": "The right-side leaning target was presented",
        "8": "The left-side leaning target was presented",
        "13": "The right mouse button was pressed",
        "12": "The left mouse button was pressed"
    }
  },
  "feedback":{
    "Description": "The visual feedback the users recived to see if their resposne correct or not.",
    "Levels":{
      "smiley_face": "A green smiley face would show at the middle of the screen for 400 ms",
      "sad_face": "A red sad face would show at the middle of the screen for 400 ms"
    },
    "HED":{
      "smiley_face": "(Visual-presentation, (Feedback, True), (Foreground-view, (Center-of, Computer-screen), (Circle, Green)), Label/Smiley-face)",
      "sad_face": "(Visual-presentation, (Feedback, False), (Foreground-view, (Center-of, Computer-screen), (Circle, Red)), Label/Sad-face)"
    }
  },
  "hed_defs":{
    "Description": "Metadata dictionary for defining the stimulus",
    "HED":{
      "contrastChange": "(Definition/contrastChange,((Visual-presentation, (Background-view, Gray), (Foreground-view, (Dots, Item-count/1, White, (Center-of, Computer-screen)))), Recording))",
      "right-target": "(Definition/right-target, (Visual-presentation, (Background-view, Gray), (Foreground-view, ((Circle, (Center-of, Computer-screen), (Radius, Angle/6 degree), (Pattern, Rightward, Label/Grating), (Increasing, Luminance-Contrast)), (Circle, (Center-of, Computer-screen), (Radius, Angle/6 degree), (Pattern, Leftward, Label/Grating), (Decreasing, Luminance-Contrast))))))",
      "right-comeback": "(Definition/right-comeback, (Visual-presentation, (Background-view, Gray), (Foreground-view, ((Circle, (Center-of, Computer-screen), (Radius, Angle/6 degree), (Pattern, Rightward, Label/Grating), (Decreasing, Luminance-Contrast)), (Circle, (Center-of, Computer-screen), (Radius, Angle/6 degree), (Pattern, Leftward, Label/Grating), (Increasing, Luminance-Contrast))))))",
      "left-target": "(Definition/left-target, (Visual-presentation, (Background-view, Gray), (Foreground-view, ((Circle, (Center-of, Computer-screen), (Radius, Angle/6 degree), (Pattern, Leftward, Label/Grating), (Increasing, Luminance-Contrast)), (Circle, (Center-of, Computer-screen), (Radius, Angle/6 degree), (Pattern, Rightward,
```

```
Label/Grating), (Decreasing, Luminance-Contrast))))))",
      "left-comeback": "(Definition/left-comeback, (Visual-presentation, (Background-view, Gray), (Foreground-view, ((Circle, (Center-of, Computer-screen), (Radius, Angle/6 degree), (Pattern, Leftward, Label/Grating), (Decreasing, Luminance-Contrast)), (Circle, (Center-of, Computer-screen), (Radius, Angle/6 degree), (Pattern, Rightward, Label/Grating), (Increasing, Luminance-Contrast))))))"
    }
  }
}
```
