## Supplementary material S5 for "HBN-EEG: The FAIR implementation of the Healthy Brain Network (HBN) electroencephalography dataset"

```
{
  "onset": {
    "LongName": "Event onset",
    "Description": "Position (latency) of event marker in seconds relative to the start.",
    "Units": "seconds"
  },
  "duration": {
    "Description": "Duration of the event, set typically to n/a. Instead, the duration of the event is determined by the onset/offset flags or the next event.",
    "Units": "seconds"
  },
  "sample": {
    "Description": "Temporal position (latency) of event marker relative to the start, in data frames."
  },
  "value": {
    "Description": "The event marker value (type).",
    "Levels": {
        "seqLearning_start": "Sequence Learning task started",
        "seqLearning_stop": "Sequence Learning task stopped",
        "learningBlock_1": "Start of the first block of sequence learning",
        "learningBlock_2": "Start of the second block of sequence learning",
        "learningBlock_3": "Start of the thrid block of sequence learning",
        "learningBlock_4": "Start of the forth block of sequence learning",
        "learningBlock_5": "Start of the fifth block of sequence learning",
        "dot_no1_ON": "Dot number 1 turned black",
        "dot_no1_OFF": "Dot number 1 turned white",
        "dot_no2_ON": "Dot number 2 turned black",
        "dot_no2_OFF": "Dot number 2 turned white",
        "dot_no3_ON": "Dot number 3 turned black",
        "dot_no3_OFF": "Dot number 3 turned white",
        "dot_no4_ON": "Dot number 4 turned black",
        "dot_no4_OFF": "Dot number 4 turned white",
        "dot_no5_ON": "Dot number 5 turned black",
        "dot_no5_OFF": "Dot number 5 turned white",
        "dot_no6_ON": "Dot number 6 turned black",
        "dot_no6_OFF": "Dot number 6 turned white",
        "dot_no7_ON": "Dot number 7 turned black",
        "dot_no7_OFF": "Dot number 7 turned white",
        "dot_no8_ON": "Dot number 8 turned black",
        "dot_no8_OFF": "Dot number 8 turned white"
    },
    "HED":{
      "seqLearning_start": "(Duration/1 s, (Visual-presentation, (Background-view,
```

```
Gray), (Foreground-view, (White, (Sentence, Label/THE-TASK-STARTS-NOW))))), (Visual-presentation, (Background-view, Gray), (Foreground-view, (White, (Sentence, Label/THE-TASK-STARTS-NOW)))), Recording",
      "learningBlock_1": "(Def/learningBlock, Onset), (Delay/2 s, ((White, Pattern, Circle), {target_count}))",
      "learningBlock_2": "(Def/learningBlock, Onset), (Delay/2 s, ((White, Pattern, Circle), {target_count}))",
      "learningBlock_3": "(Def/learningBlock, Onset), (Delay/2 s, ((White, Pattern, Circle), {target_count}))",
      "learningBlock_4": "(Def/learningBlock, Onset), (Delay/2 s, ((White, Pattern, Circle), {target_count}))",
      "learningBlock_5": "(Def/learningBlock, Onset), (Delay/2 s, ((White, Pattern, Circle), {target_count}))",
      "dot_no1_ON": "(Def/dot-no-1, Onset)",
      "dot_no1_OFF": "(Def/dot-no-1, Offset)",
      "dot_no2_ON": "(Def/dot-no-1, Onset)",
      "dot_no2_OFF": "(Def/dot-no-1, Offset)",
      "dot_no3_ON": "(Def/dot-no-1, Onset)",
      "dot_no3_OFF": "(Def/dot-no-1, Offset)",
      "dot_no4_ON": "(Def/dot-no-1, Onset)",
      "dot_no4_OFF": "(Def/dot-no-1, Offset)",
      "dot_no5_ON": "(Def/dot-no-1, Onset)",
      "dot_no5_OFF": "(Def/dot-no-1, Offset)",
      "dot_no6_ON": "(Def/dot-no-1, Onset)",
      "dot_no6_OFF": "(Def/dot-no-1, Offset)",
      "dot_no7_ON": "(Def/dot-no-1, Onset)",
      "dot_no7_OFF": "(Def/dot-no-1, Offset)",
      "dot_no8_ON": "(Def/dot-no-1, Onset)",
      "dot_no8_OFF": "(Def/dot-no-1, Offset)"
    }
  },
  "event_code": {
    "Description": "The original code used during data collection to indicate an event marker",
    "Levels": {
        "91": "Sequence Learning task started",
        "50": "Sequence Learning task stopped",
        "31": "Start of the first block of sequence learning",
        "32": "Start of the second block of sequence learning",
        "33": "Start of the thrid block of sequence learning",
        "34": "Start of the forth block of sequence learning",
        "35": "Start of the fifth block of sequence learning",
        "11": "Dot number 1 turned black",
        "21": "Dot number 1 turned white",
        "12": "Dot number 2 turned black",
        "22": "Dot number 2 turned white",
```

```
          "13": "Dot number 3 turned black",
          "23": "Dot number 3 turned white",
          "14": "Dot number 4 turned black",
          "24": "Dot number 4 turned white",
          "15": "Dot number 5 turned black",
          "25": "Dot number 5 turned white",
          "16": "Dot number 6 turned black",
          "26": "Dot number 6 turned white",
          "17": "Dot number 7 turned black",
          "27": "Dot number 7 turned white",
          "18": "Dot number 8 turned black",
          "28": "Dot number 8 turned white"
      }
  },
  "user_answer":{
    "Description": "The sequence that subject responded when asked to repeat the shown sequence",
    "HED": "Agent-action, (Participant-response, Label/#)"
  },
  "correct_answer":{
    "Description": "The correct sequence (correct sequence is the same for all blocks)",
    "HED": "Label/#"
  },
  "target_count":{
    "Description": "The number of target circles in the sequence",
    "Levels": {
      "6": "The sequence has 6 targets",
      "8": "The sequence has 8 targets"
    },
    "HED": {
      "6": "(Item-count/6)",
      "8": "(Item-count/8)"
    }
  },
  "hed_defs":{
    "Description": "Metadata dictionary for defining the stimulus",
    "HED":{
      "learningBlock": "(Definition/learningBlock, (Visual-presentation, (Background-view, Gray)))",
      "dot-no-1_def": "(Definition/dot-no-1, ((Visual-presentation, (White, Circle, (Horizontally-oriented, Angle/0 degree), Label/Filled))))",
      "dot-no-2_def": "(Definition/dot-no-2, ((Visual-presentation, (White, Circle, (Horizontally-oriented, Angle/-45 degree), Label/Filled))))",
      "dot-no-3_def": "(Definition/dot-no-3, ((Visual-presentation, (White, Circle,
```

```
(Horizontally-oriented, Angle/-90 degree), Label/Filled))))",
      "dot-no-4_def": "(Definition/dot-no-4, ((Visual-presentation, (White, Circle, (Horizontally-oriented, Angle/-135 degree), Label/Filled))))",
      "dot-no-5_def": "(Definition/dot-no-5, ((Visual-presentation, (White, Circle, (Horizontally-oriented, Angle/-180 degree), Label/Filled))))",
      "dot-no-6_def": "(Definition/dot-no-6, ((Visual-presentation, (White, Circle, (Horizontally-oriented, Angle/-225 degree), Label/Filled))))",
      "dot-no-7_def": "(Definition/dot-no-7, ((Visual-presentation, (White, Circle, (Horizontally-oriented, Angle/-270 degree), Label/Filled))))",
      "dot-no-8_def": "(Definition/dot-no-8, ((Visual-presentation, (White, Circle, (Horizontally-oriented, Angle/-315 degree), Label/Filled))))"

    }
  }
}
```
