## Supplementary material S6 for "HBN-EEG: The FAIR implementation of the Healthy Brain Network (HBN) electroencephalography dataset"

```
{
  "onset": {
    "LongName": "Event onset",
    "Description": "Position (latency) of event marker in seconds relative to the start.",
    "Units": "seconds"
  },
  "duration": {
    "Description": "Duration of the event, set typically to n/a. Instead, the duration of the event is determined by the onset/offset flags or the next event.",
    "Units": "seconds"
  },
  "sample": {
    "Description": "Temporal position (latency) of event marker relative to the start, in data frames."
  },
  "value": {
    "Description": "The event marker value (type).",
    "Levels": {
      "symbolSearch_start": "Symbol Search task started",
      "newPage": "New page of symbol search queries was shown.",
      "trialResponse": "Subject provided a resposne"
    },
    "HED": {
      "symbolSearch_start": "(Def/symbolSearch-start, Onset)",
      "newPage": "(Def/new-page, Onset)",
      "trialResponse": "Agent-action, Participant-response, (Press, (Mouse-button, (Left-side-of, Computer-mouse)), {user_answer})"
    }
  },
  "event_code": {
  "Description": "The original code used during data collection to indicate an event marker",
    "Levels": {
      "92": "Symbol Search task started",
      "20": "New page of symbol search queries was shown",
      "14": "Subject provided a resposne"
    }
  },
  "user_answer":{
    "Description": "content of user's response",
    "Levels": {
      "0": "False, symbol was not found",
      "1": "True, symbol was found"
    },
```

```
    "HED": {
      "0": "(Label/response-value, False)",
      "1": "(Label/response-value, True)"
    }
  },
  "correct_answer":{
    "Description": "the correct response",
    "Levels": {
      "0": "Symbol was not present in the search query",
      "1": "Symbol was present in the search query"
    },
    "HED": {
      "0": "(Label/correct-value, False)",
      "1": "(Label/correct-value, True)"
    }
  },
  "hed_defs":{
    "Description": "Metadata dictionary for defining the stimulus",
    "HED":{
      "symbolSearch-start_def": "(Definition/symbolSearch-start,((Visual-presentation, (Background-view, White)), Recording))",
      "new-page_def": "(Definition/new-page, (Visual-presentation, (Questionnaire, (Label/Symbol, Item-count/7), Item-count/15)))"
    }
  }}
```
